# The role of *Cyp19a1* in female pathway of a freshwater turtle (*Mauremys reevesii*) with temperature-dependent sex determination

**DOI:** 10.1101/2021.05.17.444498

**Authors:** Peng-fei Wu, Xi-feng Wang, Fei Gao, Wei-guo Du

## Abstract

The molecular mechanism of temperature-dependent sex determination (TSD) in reptiles has been drawn great interest from biologists for several decades. However, which genetic factors are essential for TSD remain elusive, especially for the female sex determination process. *Cyp19a1*, encodes an enzyme of aromatase catalyzing the conversion of testosterone to estrogen, has been confirmed to modulate steroid hormones involved in the sexual differentiation of many species, but whether it has a critical role in determining the gonadal sexual fate in TSD is still to be elucidated. Here, we identified that *Cyp19a1* expression exhibited a temperature-dependent, sexually dimorphic expression pattern, preceding gonadal sex differentiation in a TSD turtle *Mauremys reevesii. Cyp19a1* expression in gonads increased dramatically when embryos developed at high female-producing temperatures (FPT), but were extremely low throughout embryogenesis at low male-producing temperatures (MPT). *Cyp19a1* expression increased rapidly in response to the temperature shift from MPT to FPT in developing gonads. The sexual phenotype of turtles was successfully reversed by aromatase inhibitor treatment at FPT, and by estrogen treatment at MPT, accompanied with the rapid upregulation of *Cyp19a1*. These results demonstrate that *Cyp19a1* is essential for the female sex determination process in M. reevesii, indicating its vital role in the female pathway of TSD.

## 1 Introduction

Two sexes universally exist in many animals, and a diverse of mechanisms that determine sex have evolved across different lineages. In vertebrates, sex determination mainly divided into two types: genotype sex determination (GSD) and environmental-dependent sex determination (ESD) (Valenzuela and Lance, 2004). Temperature-dependent sex determination (TSD) is a notable ESD existing in some reptiles, in which the incubation temperature of developing embryos determines the gonadal sex (Charnier, 1966; Ferguson and Joanen, 1982; Pieau et al., 1999).

In reptiles, the molecular mechanisms underlying TSD have undergone several decades of research. Early studies mainly focused on hormone-dependent mechanisms, and clone and/or expression of some classical sex-related genes involved in GSD system (e.g., *Wt1; Dmrt1; Amh*) in TSD species (Gutzke and Bull, 1986; Wibbels and Crews, 1992; Spotila and Hall, 1998; Pieau et al., 1999; Gabriel et al., 2001). Later studies demonstrated that several of those genes exhibited temperature-dependent expression patterns during thermo-sensitive period (TSP), prior to gonadal sex differentiation (Shoemaker et al., 2007a; Shoemaker-Daly et al., 2010; Matsumoto and Crews, 2012). Recently, transcriptomes of TSD taxa identified a great number of candidate sex-determining genes in TSD reptiles (Czerwinski et al., 2016; Yatsu et al., 2016; Radhakrishnan et al., 2017). In addition, genetic manipulation techniques (e.g., RNA interference) were introduced to identify the function of sex-related genes in sex determination of reptiles (Shoemaker-Daly et al., 2010; Sifuentes-Romero et al., 2013; Ge et al., 2017). For example, *Dmrt1* determines the male fate in a TSD turtle *Trachemys scripta* through the *Temperature-Ca2+-pSTAT3-Kdm6b-Dmrt1* pathway, revealing a direct genetic link between epigenetic mechanism and TSD (Ge et al., 2017; Ge et al., 2018; Weber et al., 2020). Despite several decades of research on TSD mechanisms in reptiles, the role and function of key sex-related genes in female sex determination process remains unverified using robust molecular evidences including gene expression and genetic manipulation.

*Cyp19a1* (Cytochrome P450 Family 19 Subfamily A Member 1) encodes an endoplasmic reticulum enzyme called aromatase, catalyzing the conversion of testosterone to estrogen (Simpson et al., 1994), which is critical for estrogen synthesis. Estrogen plays a critical role in the sexually dimorphic development of anatomical, functional and behavioral characteristics that are essential for female development (Nelson and Habibi, 2013; Hamilton et al., 2014; Tokunaga et al., 2014). *Cyp19a1* has been identified to modulate steroid hormones involving in the sexual differentiation of GSD species (Elbrecht and Smith, 1992; Crews and Bergeron, 1994; Kitano et al., 2000; Sakata et al., 2005). In GSD vertebrates like medaka (*Oryzias latipes*) and chicken (*Gallus gallus*), *Cyp19a1* generally exhibits a sexually dimorphic expression pattern in female gonads at early embryonic days and robustly expresses within the cytoplasm in ovarian medullas; the gain- and loss-of-function analyses further provide strong evidence that *Cyp19a1* is both necessary and sufficient for ovarian differentiation (Lambeth et al., 2013; Mawaribuchi et al., 2014; Nakamoto et al., 2018; Jin et al., 2020). Although *Cyp19a1* is not required for the fetal sexual differentiation in mouse (*Mus musculus*), it is essential for granulosa cell phenotype maintaining and follicle formation in postnatal mammalian ovaries (ERICKSO et al., 1979; Britt et al., 2001; Meseke et al., 2018).

Given its vital role in GSD system, *Cyp19a1* has long been deemed to be involved in the female pathway of TSD (Ramsey et al., 2007; Fernandino et al., 2008; Parrott et al., 2014; Tang et al., 2017). Previous studies demonstrated that *Cyp19a1* transcripts were detected in adrenal-kidney-gonad complexes (AKGs) after the TSP of sex determination in the American alligator (*Alligator mississippiensis*) and several TSD turtles, denying its role as an initial trigger for sex determination (Gabriel et al., 2001; Murdock et al., 2003; Valenzuela and Shikano, 2007). In contrast, recent transcriptomic data showed that *Cyp19a1* exhibited a female-specific expression pattern in early embryonic gonads during the late TSP (embryonic developmental stage 19) in several TSD turtles (Czerwinski et al., 2016; Radhakrishnan et al., 2017). Therefore, the role of *Cyp19a1* in TSD remains unsolved in reptiles and may differ among species. Moreover, these previous studies merely provide the mRNA expression of *Cyp19a1* in embryonic gonads or even AKGs, which is not enough for making an unequivocal conclusion of *Cyp19a1* function. Further studies involving multi-level evidences (e.g., gene expression pattern, gene thermo-sensitivity, hormone/genetic manipulation analysis) in embryonic gonads are needed to identify the function of this gene.

In our previous study, we have identified that *Cyp19a1* exhibited a differential expression pattern in the AKGs between male-producing temperature (MPT) and female-producing temperature (FPT) in the freshwater turtle *Mauremys reevesii*, a TSD species (Tang et al., 2017). In this study, we firstly identified the expression pattern of *Cyp19a1* by sampled embryonic gonads at different developmental embryonic stages at FPT and MPT, and further examined the thermo-sensitivity of *Cyp19a1* via temperature shifts incubation experiments in this turtle species. To verify the role of *Cyp19a1* in inducing female differentiation, we administered estrogen to MPT embryos to induce male-to-female sex reversal and detected the *Cyp19a1* expression. Furthermore, we examined the necessity of *Cyp19a1* in female determination process through handling aromatase inhibitor to FPT embryos. Collectively, our findings revealed that *Cyp19a1* played a vital role in the female sex determination pathway of a TSD turtle, which provided new insight into our understanding of female sex-determining pathways in TSD reptiles.

## 2 RESULTS

### 2.1 Characterization of *Cyp19a1* gene in *M. reevesii*

In *M. reevesii*, the complete cDNA sequence of *Cyp19a1* was 1995 base pairs (bp), with a 126 bp 5’ untranslated region (UTR), a 355 bp 3’ UTR and an open reading frame (ORF) of 1515 bp that encoded a protein of 504 amino acids (Fig. S1A). The deduced amino acid sequence of *M. reevesii Cyp19a1* shared 84.9%, 84.4%, 75.1%, and 53.5% identity with that of chicken, lizard, human, and zebrafish, respectively (Fig. S1B), indicating that *M. reevesii Cyp19a1* was evolutionary more closely related to chicken, lizard and human than to fish (Fig. S1C).

*Cyp19a1* mRNA was abundantly expressed in ovary, but not in testis of adult turtles (Fig. S2A). The adult testis had a dense medulla with seminiferous cords, while ovary showed a developing follicular system (Fig. S2B).

### 2.2 Sexually dimorphic expression of *Cyp19a1* in early embryonic gonads of *M. reevesii*

The *Cyp19a1* transcripts started to express in FPT gonads at embryonic developmental stage of 16, which occurred at early TSP, and clearly prior to gonadal sexual differentiation. *Cyp19a1* expression increased dramatically as embryos developed: the transcripts of *Cyp19a1* in FPT gonads was only dozens of times of MPT gonads at developmental stage of 16, but was thousands of times or even higher at developmental stage of 19 or 25. In contrast, MPT gonads exhibited extremely low expression of *Cyp19a1* throughout embryogenesis (Fig. 1).

**Fig. 1.**
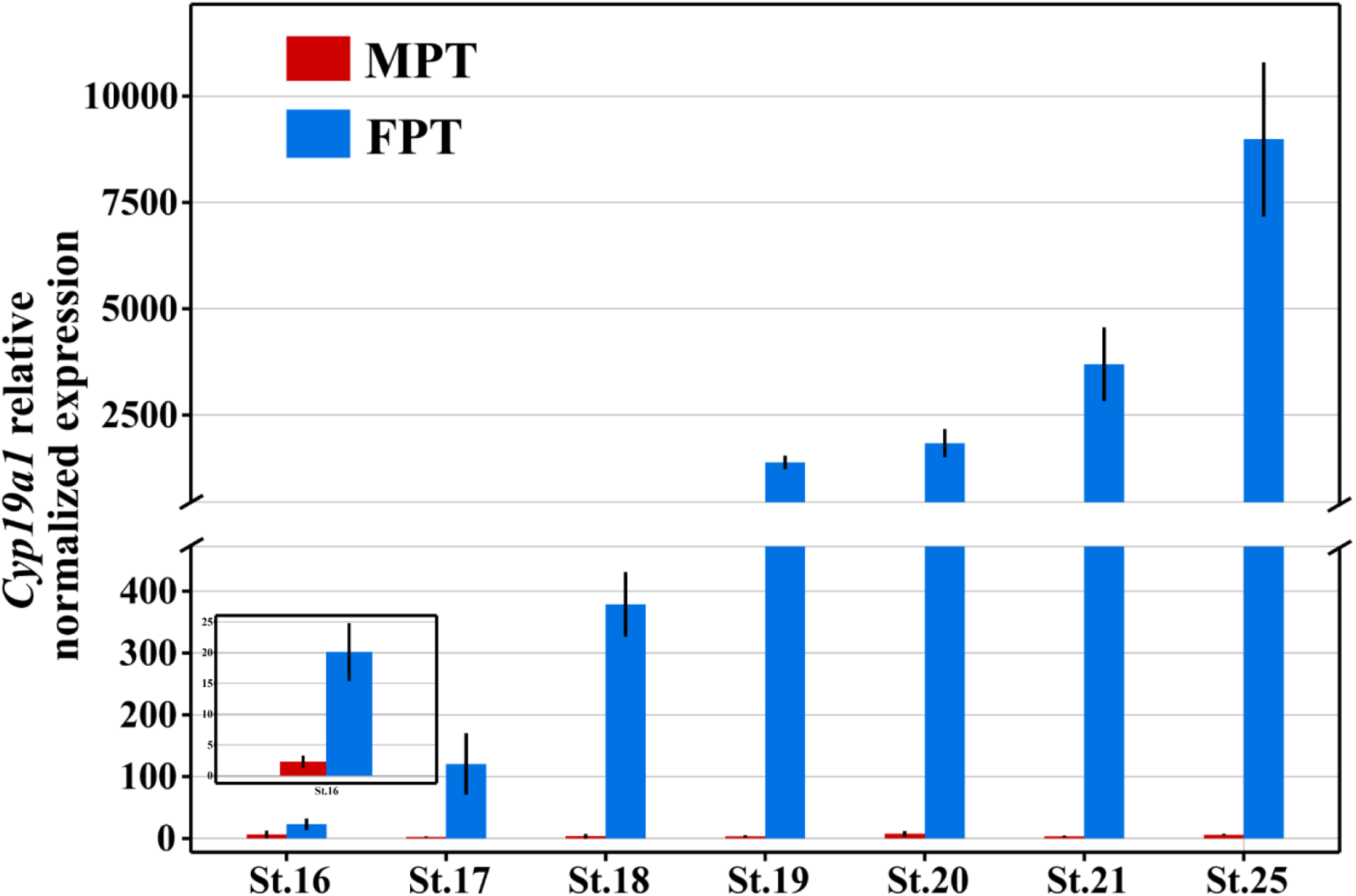
The sexually dimorphic expression pattern of *Cyp19a1* in *M. reevesii*. The mRNA expression of *Cyp19a1* in gonads of different stages (16-25) at MPT (26°C) and FPT (32°C) determined by qRT-PCR analysis; β-actin was used as a reference gene. *Cyp19a1* exhibited a highly FPT-specific expression pattern in early embryonic gonads. Data are mean±s.d.; n≥3.\

### 2.3 Thermo-sensitivity of *Cyp19a1* in *M. reevesii*

The expression level of *Cyp19a1* in turtle gonads obviously decreased with reducing temperatures from 31°C to 27°C at both developmental stage of 18 (during TSP) and 21 (after TSP) (Fig. 2A, B). In addition, *Cyp19a1* expression responded rapidly to the temperature shift in developing gonads when embryos were transferred from MPT to FPT at stage 16. *Cyp19a1* increased slightly above the MPT-typical level at early stage 17 and reached significantly high levels at the subsequent stages. Interestingly, *Cyp19a1* expression showed a time lag in response to the temperature shift in gonads transferred from FPT to MPT, continuously upregulating until stage 19 and then decreasing to the MPT-typical level (Fig. 2C).

**Fig. 2.**
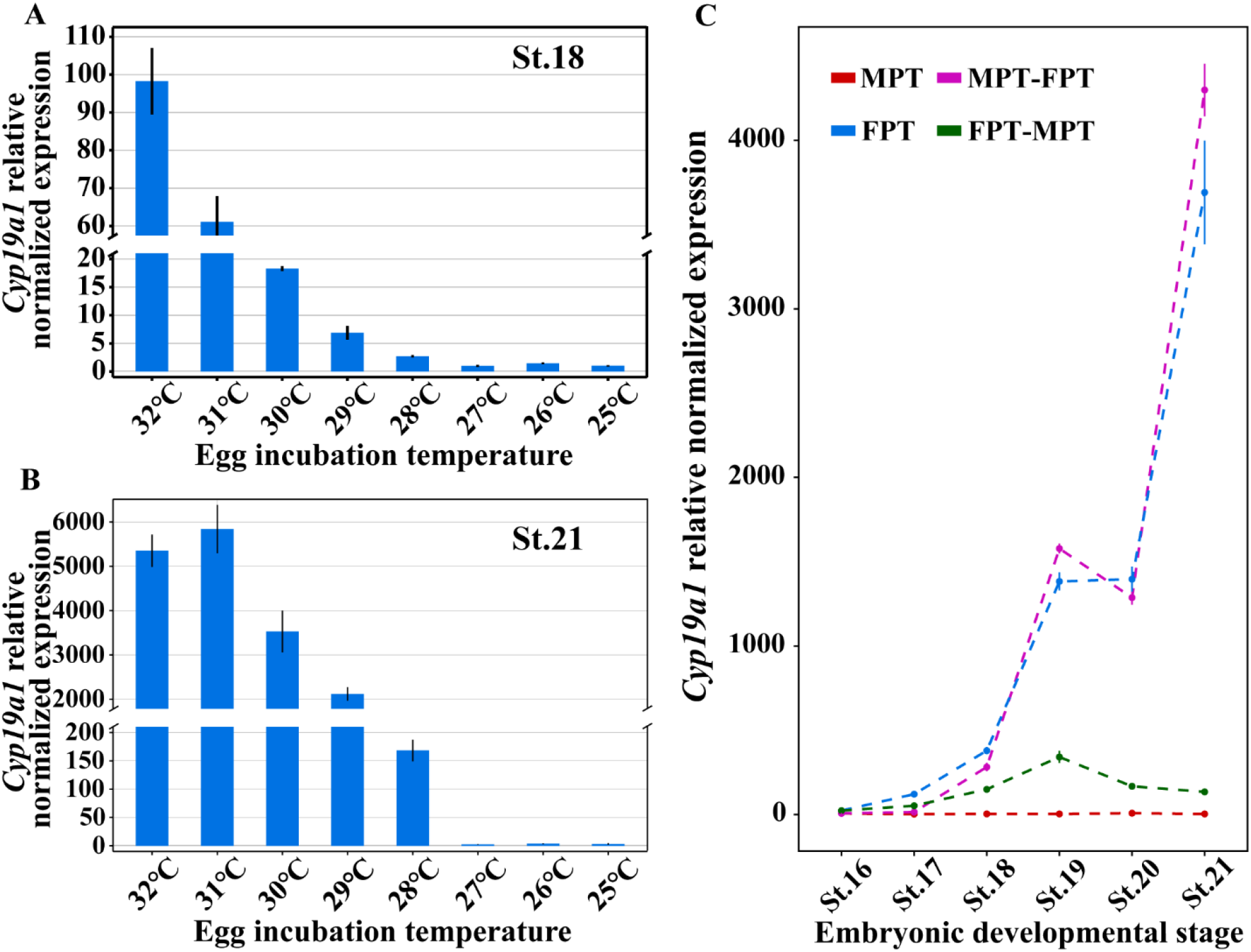
Thermo-sensitivity of *Cyp19a1* in *M. reevesii*. (A, B) qRT-PCR analysis of *Cyp19a1* in gonads at stage 18 or stage 21 of gradient temperatures. The mRNA expression of *Cyp19a1* in embryonic gonads exhibited temperature dependence, no matter during the TSP (A) or after (B). (C) Time-course response of *Cyp19a1* expression to temperature shifts from either MPT→FPT or FPT→ MPT. Embryos were shifted at stage 16, and gonads were dissected for qRT-PCR analysis at stages 16-21. *Cyp19a1* expression in gonads with temperature shifts from MPT→FPT responded rapidly to the new temperatures, with rapid expression changes occurring at stage 17. In the opposite shift (FPT→MPT), *Cyp19a1* expression continuously increased a little until stage 19, and then decreased towards the MPT-typical level. Data are mean±s.d.; n≥3. P<0.05.

### 2.4 Upregulation of *Cyp19a1* in feminized MPT gonads during male-to-female sex reversal in *M. reevesii*

In MPT embryos with estrogen treatment, the gonads exhibited female-like morphology, with a degenerated medulla and a thickened outer cortex at stage 25 (Fig. 3A-F’). Meanwhile, a significant upregulation in *Foxl2* and *Rspo1*, and downregulation of *Dmrt1* and *Sox9* were detected in feminized MPT embryos at stage 25 (Fig. 3G). *Sox9* protein was highly expressed in the medulla of MPT gonads, but decreased dramatically in feminized MPT gonads induced by estrogen at either developmental stage of 21 or 25 (Fig. 3H-J”, Fig. S3A-C”). Correspondingly, germ cells in MPT gonads with estrogen treatments mainly enriched in the developed outer cortex with few germ cells localized in the medulla (Fig. 3L-L”, Fig. S3E-E”), exhibiting a female-like distribution pattern (cortical localization of germ cells) rather than a male pattern (medullary cord distribution of germ cells) (Fig. 3K-K”, M-M”, Fig. S3D-D”, F-F”). Moreover, *Cyp19a1* transcripts responded rapidly to estrogen treatment and were significantly upregulated from stage 17 onwards (Fig. 3N).

**Fig. 3.**
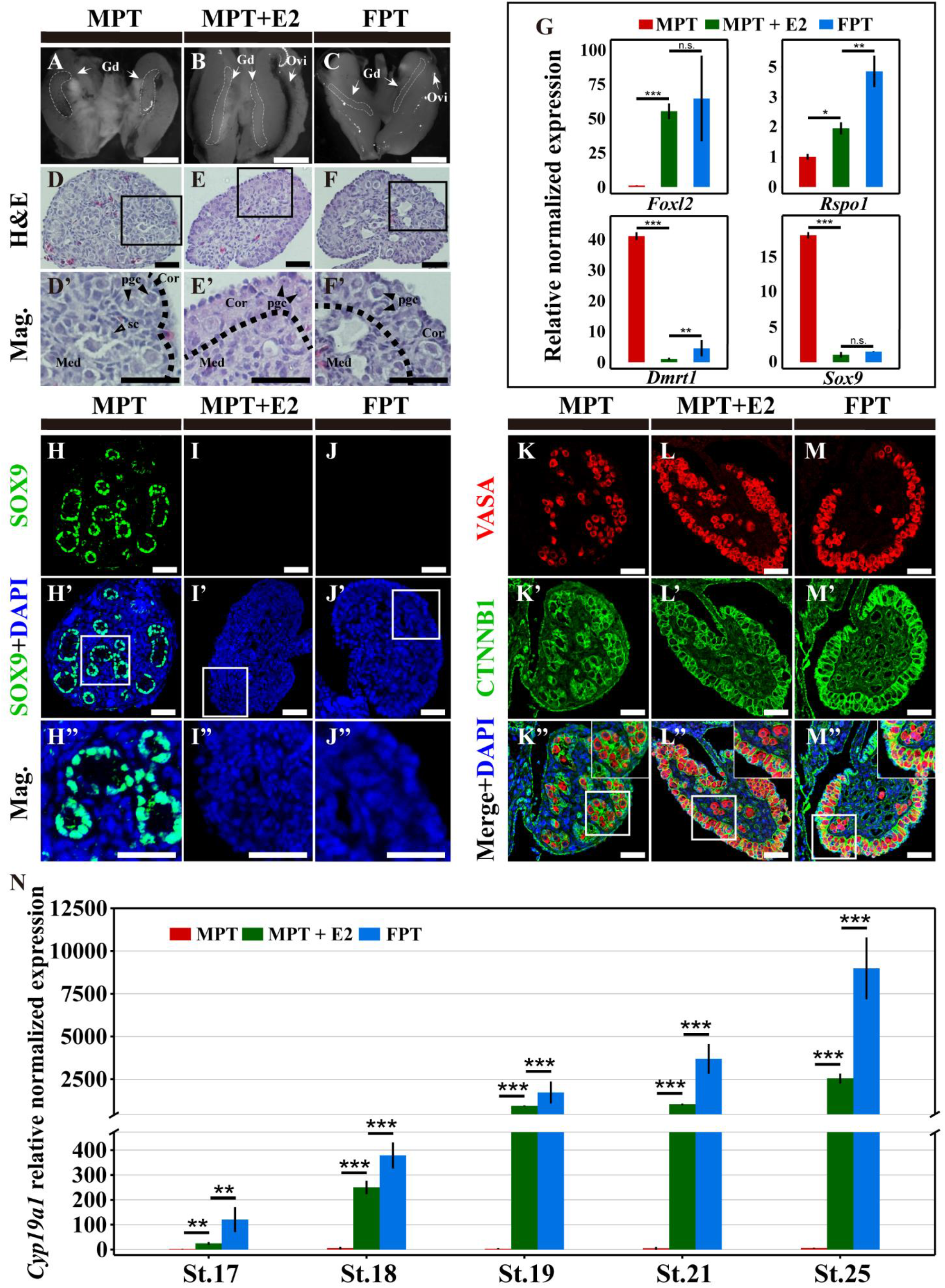
The upregulation of *Cyp19a1* in feminized MPT gonads during male-to-female sex reversal. (A-C) Gonads (outlined by white dotted lines) on top of mesonephros of MPT, feminized MPT and FPT embryos at stage 25. Gd, gonad; Ovi, oviduct. Scale bars: 1 mm. (D-F) H&E staining of gonadal sections from MPT, feminized MPT and FPT embryos at stage 25 showed that the feminized MPT embryonic gonads had a degenerated medulla and a thickened outer cortex. The dashed line indicates the border between medulla and cortex. pgc, primordial germ cells; sc, seminiferous cord; Cor, cortex; Med, medulla; Scale bars: 50 μm. (G) qRT-PCR analysis of *Foxl2, Rspo1*, *Dmrt1* and *Sox9* in gonads from MPT, MPT with estrogen treatment and FPT embryos at stage 25. Significant upregulation of *Foxl2* and *Rspo1* expression and strong downregulation of *Dmrt1* and *Sox9* expression were observed in stage 25 MPT gonads with estrogen treatment. (H-J”) Protein localization of *Sox9* in MPT, feminized MPT and FPT gonadal sections at stage 25. *Sox9* expression was totally lost in feminized MPT gonads. Scale bars: 50 μ m. (K-M′) A female-typical distribution of germ cells was observed in MPT gonads with estrogen treatment at stage 25, determined by *Vasa* and *β-catenin* immunostaining. Scale bars: 50 μ m. (N) qRT-PCR analysis of *Cyp19a1* in gonads from MPT, feminized MPT and FPT embryos at stages 17, 18, 19, 21 and 25. *Cyp19a1* exhibited a rapid response to estrogen treatment and was rapidly upregulated at stage 17. Data are mean±s.d.; *P<0.05; **P<0.01; ***P<0.001; n≥3.

### 2.5 Masculinization of FPT embryos following aromatase inhibitor treatment in *M. reevesii*

In FPT embryos treated with aromatase inhibitor, gonads became shortened and vascularized, exhibiting male-like morphology, characterized by a dense medulla with a number of primordial germ cells and a degenerated cortex (Fig. 4A-E). *Sox9* protein expressed specifically in the nuclei of precursor Sertoli cells in MPT gonads, consistent with the FPT gonads treated by aromatase inhibitor (Fig. 4G-I”, Fig. S4A-C”). The distribution of vasa-positive germ cells in FPT gonads following aromatase inhibitor treatment displayed a male-like medulla localization pattern (Fig. 4J-L”, Fig. S4D-F”).

**Fig. 4.**
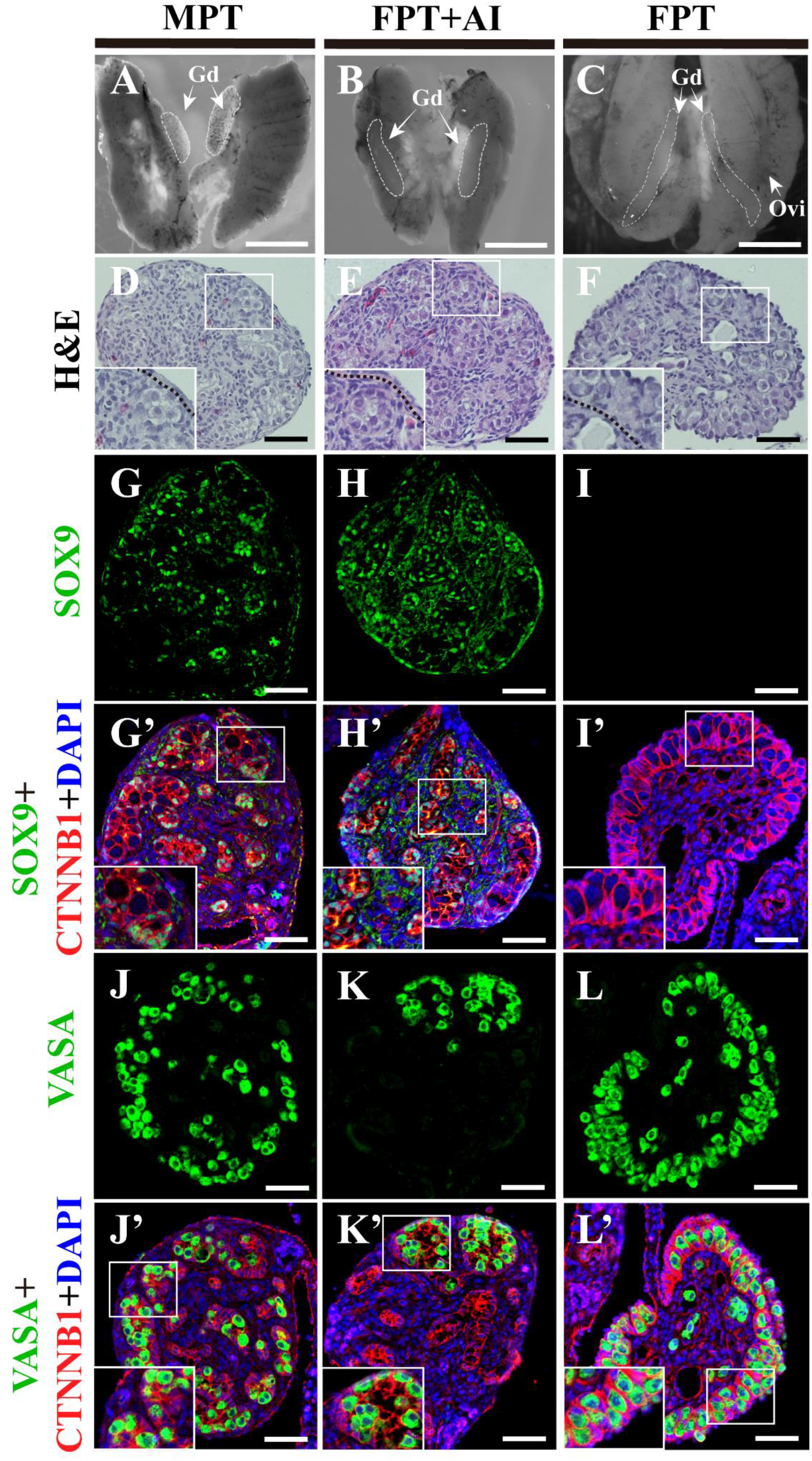
Masculinization of FPT embryos following aromatase inhibitor treatment. (A-C) Representative images of the gonad-mesonephros complexes from MPT, FPT with aromatase inhibitor treatment and FPT embryos at stage 25. Gd, gonad; Ovi, oviduct. Scale bar: 1 mm. (D-F) H&E of gonadal sections from MPT, FPT with aromatase inhibitor treatment and FPT embryos at stage 25. The FPT gonads with aromatase inhibitor treatment displayed a testis-like phenotype, characterized by a thickened outer cortex and degenerated or absent testis cords. The dashed black line indicates the border between medulla and cortex. Scale bars: 50 μm. (G-I’) Immunofluorescence of Sox9 and β-catenin in gonadal sections of MPT, FPT with aromatase inhibitor treatment and FPT embryos at stage 25. *Sox9* protein expression was robustly expressed in FPT gonads following aromatase inhibitor treatment. (J-L’) Immunofluorescence of *Vasa* in MPT, FPT with aromatase inhibitor treatment and FPT gonads at stage 25. Inhibition of aromatase in FPT gonads led to male-like distribution of germ cells within seminiferous cords. Scale bars: 50 μm.

## 3 DISCUSSION

*Cyp19a1* was thought to play an important role in the TSD female pathway, but it lacks functional analysis. Supporting this long-standing assumption of *Cyp19a1* function, we found that *Cyp19a1* expression showed a temperature-dependent, sexually dimorphic expression pattern prior to gonadal sexual differentiation (Fig.1), and was capable of responding rapidly to incubation temperature shift (Fig.2). In addition, *Cyp19a1* expression rapidly upregulated in sex-reversed gonads at MPT by estrogen treatment (Fig.3), and aromatase inhibitor induced masculinization of FPT embryos (Fig.4). These results indicated that *Cyp19a1* could determine the fate of the bipotential gonad in a TSD turtle *Mauremys reevesii*, and likely played a critical role in female determination pathway of TSD reptiles.

In previous studies, sex-related gene expression studies of TSD reptiles mainly based on two types of methods: qRT-PCR by individuals’ AKGs (inaccurate sampling) (Western et al., 2000; Gabriel et al., 2001; Valenzuela and Shikano, 2007; Tang et al., 2017), and transcriptomes by individuals’ gonads (lack of identification) (Czerwinski et al., 2016; Yatsu et al., 2016; Radhakrishnan et al., 2017). These studies demonstrated that *Cyp19a1* gene exhibited a female-specific pattern during the end of TSP or even after TSP (Gabriel et al., 2001; Valenzuela and Shikano, 2007; Czerwinski et al., 2016; Tang et al., 2017), which indicated its vital role in sexual differentiation process but denied its role as an initial trigger in TSD. For example, *Cyp19a1* transcripts merely expressed in gonads or AKGs during the late TSP (stage 19) in *Trachemys scripta* and *Chrysemys picta* (Radhakrishnan et al., 2017), or after the TSP in *Alligator mississippiensis* (Gabriel et al., 2001; Czerwinski et al., 2016). Similarly, in *M. reevesii*, early study reported that *Cyp19a1* expression in FPT AKGs was slightly above the MPT AKGs after stage 17 (Tang et al., 2017). However, AKGs could mask earlier gonadal differential expression of sex-related genes in TSD turtles (Shoemaker et al., 2007b; Valenzuela et al., 2013). To remove this potential confounding, we identified the gene expression in *M. reevesii* gonads in the present study. Interestingly, we found that the FPT-specific expression of *Cyp19a1* transcripts were detected as early as stage 16 by qRT-PCR, revealing the early female-specific expression pattern of *Cyp19a1* that precedes gonadal morphological differentiation (Fig. 1). These results strongly imply that *Cyp19a1* is an early regulator for female determination in TSD turtles and involved in the early TSD female pathway but not just sexual differentiation.

*Cyp19a1* has been previously showed temperature dependency in TSD reptiles (Gabriel et al., 2001; Murdock et al., 2003; Radhakrishnan et al., 2017). In TSD turtles, the gonad *Cyp19a1* transcripts were highly expressed when incubated in 32°C, and maintained at a low level in 26°C throughout embryonic development (Czerwinski et al., 2016; Tang et al., 2017). These studies showed a sexually dimorphic expression of *Cyp19a1*, but the details of the *Cyp19a1* expression influenced by temperature has not been investigated. In this study, *Cyp19a1* expression significantly decreased during TSP and after TSP when incubation temperatures reduced from 31 °C to 27°C with 1 °C as an interval (Fig. 2A and B), indicating its temperature dependency. Moreover, we shifted embryos from MPT to FPT or from FPT to MPT, and found that *Cyp19a1* expression rapidly increased or decreased following the change of incubation temperature (Fig. 2C). These features confirmed that *Cyp19a1* as a key gene involved in temperature-induced system. The time lag of *Cyp19a1* expression when embryos were shifted from FPT to MPT might be due to *Cyp19a1* expression was upregulated by the continuous activation of STAT3, which is crucial for TSD female pathway (Weber et al., 2020). The STAT3 is long-lived molecules that would be stable for at least 8 hours in cells and may influence the signal transduction process to regulate *Cyp19a1* expression (Siewert et al., 1999; Weber et al., 2020). Collectively, these findings confirmed the highly thermo-sensitivity of *Cyp19a1* in *M. reevesii* gonadal cells, indicating its vital role in female pathway connecting temperature and sex of TSD system.

If administered during the TSP, exogenous estrogen and its synthetase aromatase could override the temperature effect to induce the feminization of MPT embryos in TSD reptiles (Janes et al., 2007; Ramsey and Crews, 2009; Matsumoto and Crews, 2012; Warner et al., 2014; Sun et al., 2016), but the process how hormones affect sex-determining genes and further influence individuals’ sexual fate remain unclarified. In this study, we treated the embryos with estrogen at stage 16 and found that the *Cyp19a1* transcripts expression rapidly upregulated since stage 17 and the gonadal sexual phenotype of MPT embryos at stage 25 permanently feminized. There is no direct experimental evidence show that estrogen would affect the expression of *Cyp19a1* in TSD reptiles. However, it has been proposed that exogenous estrogen might redirect the gonadal trajectory by interacting with the candidate sex-determining genes like *Foxl2*, constructing a direct positive feedback regulation of *estrogen-Foxl2-Cyp19a1*-estrogen, which may promote the expression of *Cyp19a1* (Matsumoto and Crews, 2012). Therefore, it is possible that the expression pattern of *Cyp19a1* reversed by exogenous estrogen is responsible for the ultimate sex-reversal in *M. reevesii*. At the least, these results suggested that *Cyp19a1* was required for early ovarian determination in TSD.

In GSD species (e.g., chicken and Chinese soft-shelled turtle), aromatase inhibitor or *Cyp19a1* loss-of-function treatments induced a permanent female-male sex reversal, which was characterized by the formation of bilateral testis with the spermatogenesis ability and an external male phenotype (Vaillant et al., 2001; Bao et al., 2017; Jin et al., 2020). As aromatase inhibitor (Letrozole) could suppress the function of aromatase by over 98% (Lamb and Adkins, 1998; Simpson et al., 2004), aromatase inhibitor treatment is a reliable alternative method to identify the function of *Cyp19a1*. In this study, suppression of *Cyp19a1* through aromatase inhibitor treatment in *M. reevesii* indeed induced a female-to-male sex reversal, which reflected in the high expression of male-specific marker genes and the medulla localization of germ cells (Fig. 3). It’s also consistent with the previous studies that aromatase inhibitor administered during the TSP could induce the masculinization of FPT embryo in TSD reptiles (Lance and Bogart, 1992; Wibbels and Crews, 1994; Ge et al., 2017). These findings demonstrated that, like its role in chicken and Chinese soft-shelled turtle, *Cyp19a1* was necessary for female determination process in TSD turtle *M. reevesii*.

Our study provides unequivocal evidences for the critical function of *Cyp19a1* in the female pathway of turtles, but questions remain. Which factors will be the direct upstream regulator of *Cyp19a1*? The transcription factor *Foxl2*, a classical sex-determining gene (De Baere et al., 2001; Schmidt et al., 2004; Uda et al., 2004; Boulanger et al., 2014), can be one of direct upstream regulators, because the transcription of *Cyp19a1* could be directly promoted by *Foxl2* in fish, goat and human (Pannetier et al., 2006; Fleming et al., 2010; Bertho et al., 2018). Another possible upstream regulator is the promoter methylation dynamics of *Cyp19a1*, which significantly correlated with temperature change and *Cyp19a1* expression (Matsumoto et al., 2013; Parrott et al., 2014; Matsumoto et al., 2016). In addition, *NRF2*, a redox-sensitive transcription factor, is also reported directly promotes *Cyp19a1* expression during syncytiotrophoblast differentiation (Muralimanoharan et al., 2018). Therefore, which genetic factors directly regulate *Cyp19a1* expression may be the key next step to elucidate the mechanism of female pathway in TSD system.

In summary, we have demonstrated that *Cyp19a1* is a key female sex-determining gene in a fresh water turtle *M. reevesii*, which exhibited TSD. This is the first time to identify the function of *Cyp19a1* in a TSD reptile, confirming the importance of *Cyp19a1* in the female sex determination process, thereby shedding new light on the elusive TSD molecular mechanism.

## 4 MATERIALS AND METHODS

### 4.1 Sequence homology comparation and phylogenetic tree construction

The full-length coding sequence of *M. reevesii Cyp19a1* have been accessed from previous study (accession number KU821113). Alignment of deducted amino acid sequences were carried out by ClustalX and GENEDOC. Multiple amino acid alignments for the tree construction were performed using ClustalW and the phylogenetic tree was constructed using the neighbor-joining method in Mega 6.0. The accession numbers of amino acid sequences used in the phylogenetic are as follows: *Homo sapiens* (AMNP_000094.2); *Mus musculus* (NP_031836.1); *Gallus gallus* (NP_001001761.2); *Alligator mississippiensis* (XM_019477372.1); *Pelodiscus sinensis* (XP_006135137.1); *Anolis carolinensis* (XP_016852003.1); *Python bivittatus* (XP_007422249.1); *Xenopus tropicalis* (NP_001090630.1); *Andrias davidianus* (ALL29317.1); *Danio rerio* (NP_571229.3).

### 4.2 Tissue-specific expression of *Cyp19a1*

Adult fresh water turtles (*M. reevesii*) were obtained from the Hanshou Turtle Farm (Hunan, China). In order to examine the tissue-specific expression of *Cyp19a1*, we collected the heart, liver, brain, intestine, skin, kidney, lung, testis and ovary of male and female adult turtles for RT-PCR analysis. And adult testes and ovaries were separately sectioned for Hematoxylin and Eosin staining.

### 4.3 Egg collection and incubation

Freshly laid *M. reevesii* turtle eggs were still obtained from the Turtle Farm. For various incubation experiments, fertilized eggs were randomized in plastic boxes with moist vermiculite and placed in incubators at different temperatures with a water potential of −220 kPa. In this species, incubation of eggs at 26°C (MPT) generates all males, whereas incubation at 32°C (FPT) generates all females (Hou, 1985; Tang et al., 2017). For sexually dimorphic expression pattern, eggs were constantly maintained in an incubator kept at 26°C or 32°C. For temperature-shift experiments, eggs were shifted at developmental stage 16 from an incubator kept at 26°C to another kept at 32°C and vice versa. For temperature gradient incubation experiments, groups of eggs were placed into incubators held at 25°C, 26°C, 27°C, 28°C, 29°C, 30°C, 31°C and 32°C. Embryos were staged by refer to criteria established by Greenbaum (2002) in *Trachemys scripta*. We sampled gonads at stages 16, 17, 18, 19, 20, 21 and/or 25 from embryos incubated at MPT, FPT, MPT→FPT and FPT→MPT and gonads at stages 18 and 21 from embryos incubated at gradient temperature of 25 ° C-32° C for qRT-PCR analysis.

### 4.4 Exogenous estrogen and aromatase inhibitor treatments

In order to examine the role of *Cyp19a1*, a steroid estrogen (β-estradiol, E8875, Sigma) was be administered to eggs incubating at MPT (26°C), meanwhile a non-steroidal aromatase inhibitor (letrozole, PHR1540, Sigma) was administered to eggs incubating at FPT (32°C). β-estradiol or letrozole were dissolved in 95% ethanol at a concentration of 10 μg/μl, and 10 μl of the drug was applied topically to the eggshell in the region adjacent to the embryo at stage 16. Controls were treated with 10 μl of 95% ethanol. Gonad-mesonephros complexes were dissected from estrogen or aromatase inhibitor treated and control embryos at stage 25 for histology and immunohistochemistry. And gonads treated with estrogen were separated from the adjacent mesonephros at stages 17, 18, 19, 21, 25, and preserved for qRT-PCR analysis.

### 4.5 RNA extraction and qRT-PCR

Gonads from embryos in each group were harvested for RNA extraction using TRIzon Reagent (CW0580, Cwbiotech) according to a routine protocol. First-strand cDNA was synthesize using 0.5μg RNA and EasyQuick RT MasterMix (CW2019, Cwbiotech) based upon the manufacturer’s protocol. Quantitative real-time PCR reactions were performed using Roche LightCycler 480 system with a SsoFast EvaGreen Supermix (#1725201, Bio-Rad). Each sample was run in triplicate. After normalization with β-actin, relative RNA levels in samples were calculated by the comparative threshold cycle (Ct) method (Schmittgen and Livak, 2008). The sequences of primers for PCR are listed in Table S1.

### 4.6 Hematoxylin and Eosin (H&E) staining

Embryo gonad-mesonephros complexes and adult gonads were immersed in 4% paraformaldehyde (PFA) overnight at 4°C, and transferred to 70% ethanol. Tissues were weighing paper wrapped and placed in processing cassettes, dehydrated through a serial alcohol gradient, embedded in paraffin wax blocks and sectioned. Before staining, 5-μm-thick tissue sections were dewaxed in xylene, rehydrated through decreasing concentrations of ethanol, and washed in PBS. And then stained with Hematoxylin and Eosin (H&E). After staining, sections were dehydrated through increasing concentrations of ethanol and xylene, and sealed with Permount TM Mounting Medium.

### 4.7 Immunofluorescence

Gonad-mesonephros complexes from turtle embryos of indicated stages were still immersed in 4% PFA overnight at 4°C, then moved through a MeOH gradient, embedded in paraffin wax and sectioned. Paraffin sections (~5 μm) were first deparaffinized, and antigens were unmasked by microwaving sections in 10 mM/L citrate buffer, pH 6.0 (15 minutes). Sections were covered with primary antibodies and incubated overnight at 4°C or 1 hour at room temperature. The primary antibodies used in this analysis included rabbit anti-Sox9 (AB5535, Chemicon, 1:1000), rabbit anti-Vasa (ab13840, Abcam, 1:200) and mouse anti-β-catenin (C7207, Sigma, 1:250). Secondary antibodies Alexa Fluor 594 donkey anti-rabbit IgG (A21207, Invitrogen) and Alexa Fluor 594 donkey anti-mouse IgG (A21203, Invitrogen), Alexa Fluor 488 donkey anti-rabbit IgG (A21206, Invitrogen), Alexa Fluor 488 donkey anti-mouse IgG (A21202, Invitrogen), which all diluted at 1:250, were used to detect primary antibodies. Nuclei were stained with DAPI. Gonad sections were imaged by confocal microscope (A1 Plus, Nikon).

## 5 Statistical analyses

Each experiment was independently repeated at least three times. All data are expressed as the mean ± s.d. Student’s unpaired t-test was used to test significance (*P<0.05; **P<0.01; ***P<0.001; n.s., no significance).

## 6 Data availability

All data necessary for confirming the conclusions in this paper are included in this article and accompanying figures and tables.

## 7 Acknowledgements

We thank Hua Ye, Qiong Zhang, Hong-xin Xie, Chun-rong Mi, Zi-han Ding and Ming-shuo Qing for their assistance on turtle eggs collection and incubation. Research was performed under approvals from the Animal Ethics Committee at the Institute of Zoology, Chinese Academy of Sciences (IOZ14001). This work was supported by the National Natural Science Foundation of China (Grant No. 32030013, Grant No. 31821001).

## 8 Author contributions

Peng-fei Wu, Xi-feng Wang and Wei-guo Du conceived and designed the study; Peng-fei Wu and Xi-feng Wang performed the experiments; Peng-fei Wu and Xi-feng Wang analyzed data; and Peng-fei Wu, Xi-feng Wang, Fei Gao and Wei-guo Du cowrote the manuscript. All authors read and approved the manuscript. The authors declare no competing or financial interests.

## Supplementary Material

**Figure S1.**
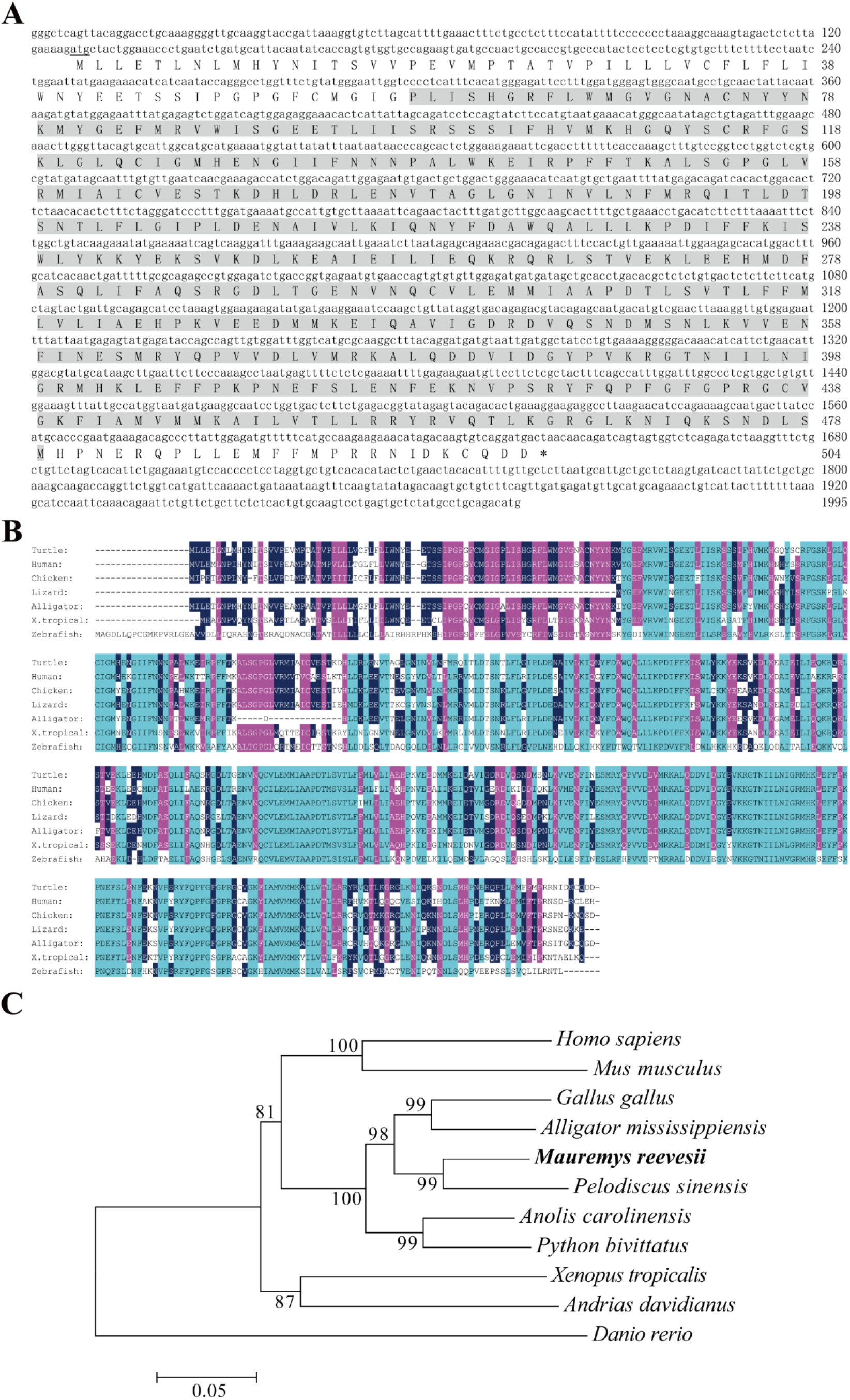
Sequence and phylogenetic analyses of *M. reevesii Cyp19a1*. (A) the complete cDNA sequence of *M. reevesii Cyp19a1* and deduced amino acid sequence. The start codon ATG was underlined, and the stop codon was indicated by an asterisk. The highly conserved cytochrome P450 domain was in shadow. (B) Alignment of amino acid sequence of *M. reevesii Cyp19a1* with those from other typical species. (C) *Cyp19a1* phylogenetic tree analysis of *M. reevesii* and other typical species based on Neighbor-Joining (N-J) method. Numbers at major branche nodes were bootstraps percentage values based on 1000 replicates. Each branch length scale in terms of genetic distance was indicated at the bottom of tree.

**Figure S2.**
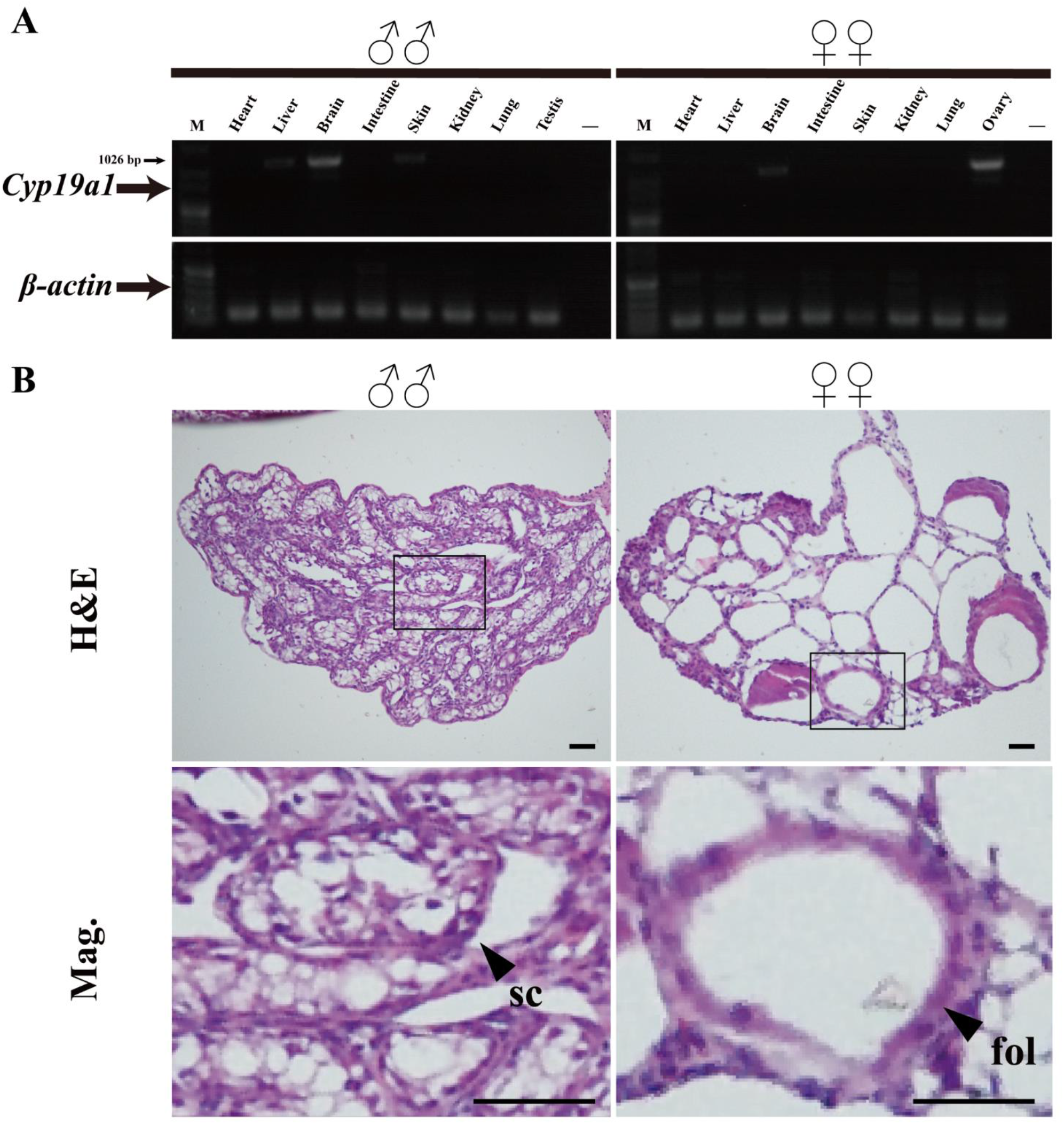
*Cyp19a1* expression and histology of testis and ovary in adult *M. reevesii*. (A) The expression of *Cyp19a1* mRNA in different tissues was analyzed by RT-PCR, which specifically high expressed in ovary instead of testis. (B) H&E staining of adult testis and ovary sections. sc, seminiferous cord; fol, follicle; scale bars are 50 μm.

**Figure S3.**
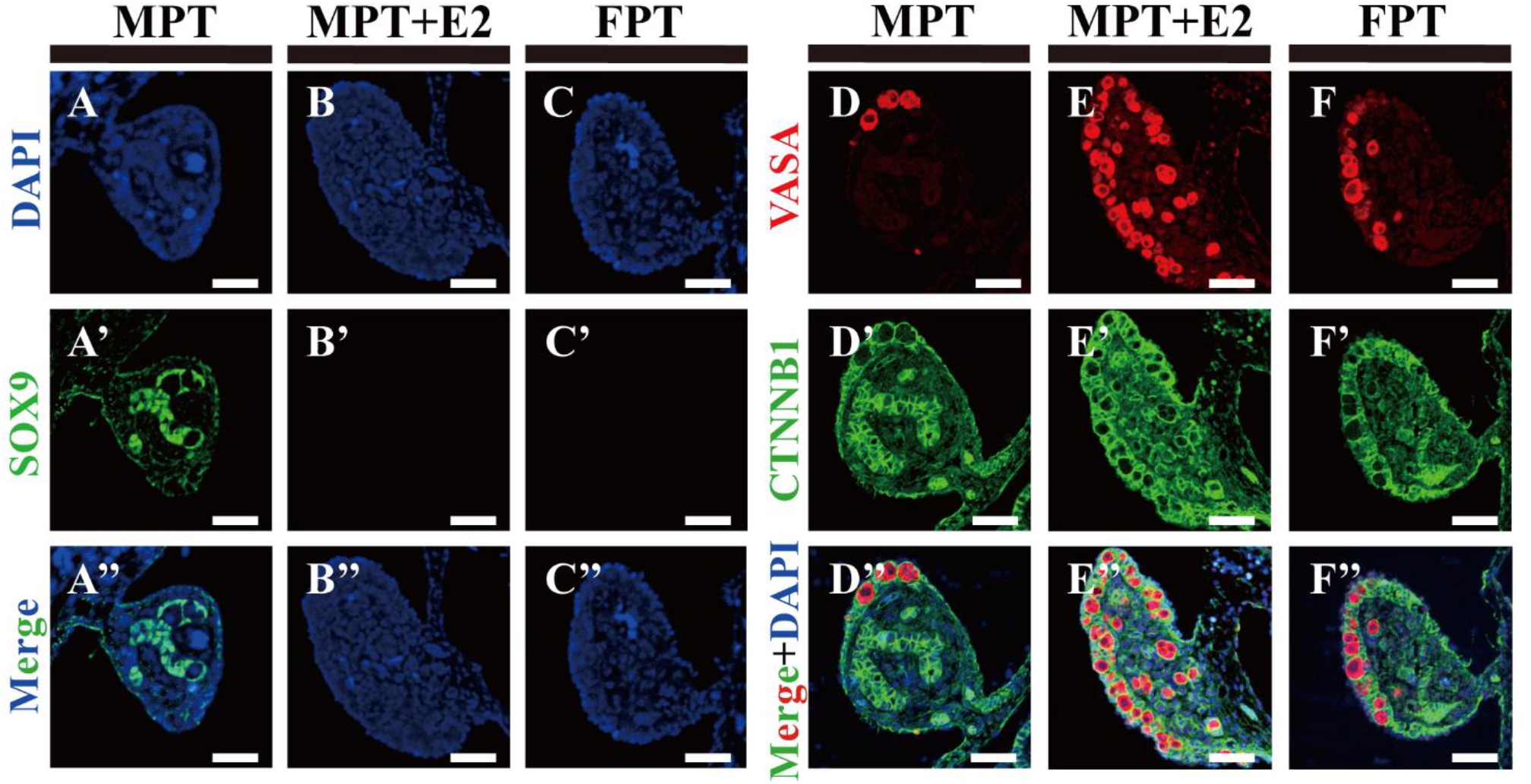
Expression of sex-specific marker genes in MPT gonads following estrogen treatment at stage 21. (A-C”) *Sox9* protein expression in gonadal sections from MPT, MPT with estrogen treatment and FPT embryos. (D-F”) The distribution of germ cells in MPT gonads following estrogen treatment, determined by *Vasa* immunostaining. Scale bars are 50 μm.

**Figure S4.**
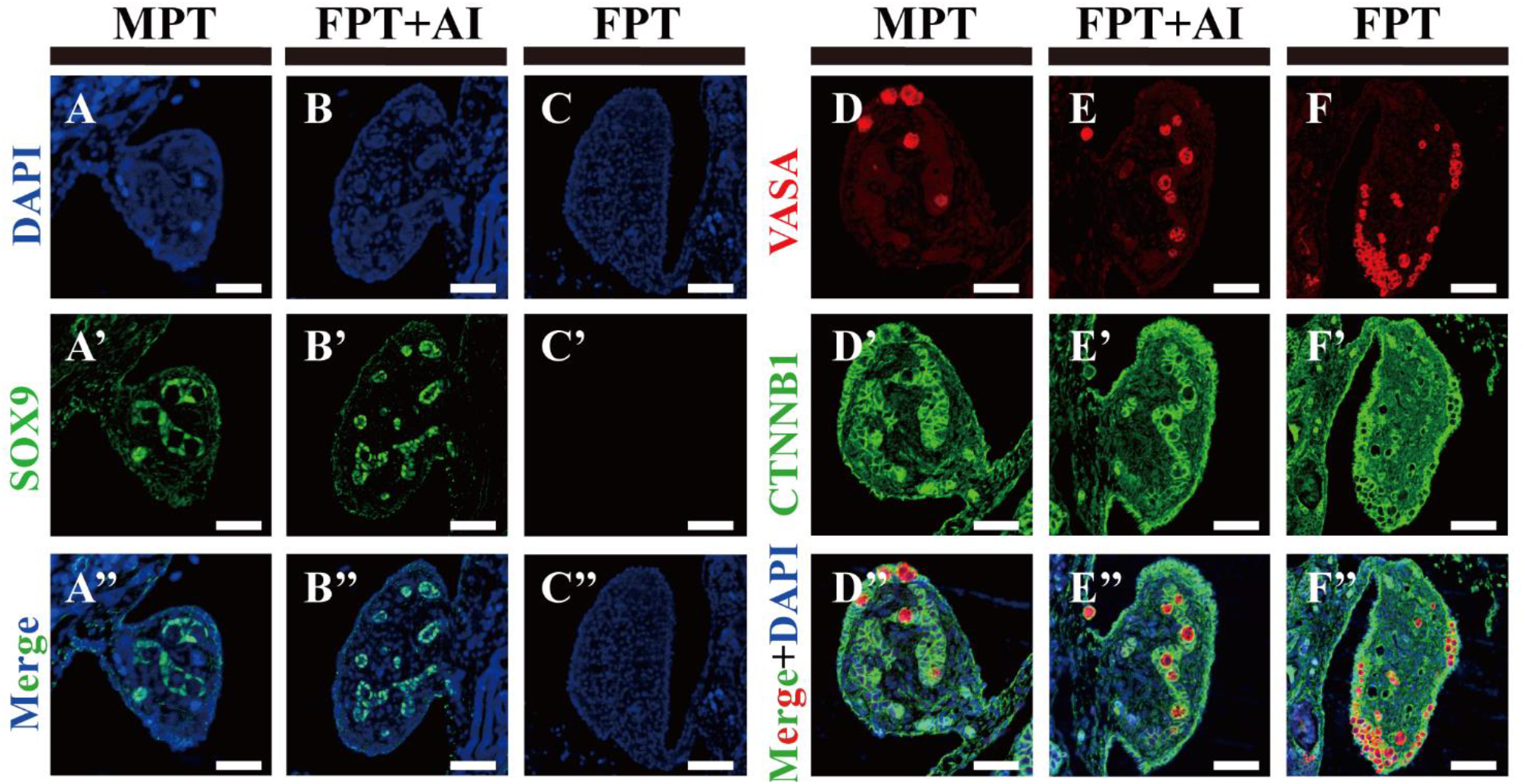
Masculinization of FPT *M. reevesii* embryos following aromatase inhibitor treatment at stage 21. (A-C”) Immunofluorescence of *Sox9* in transverse sections of MPT, FPT with aromatase inhibition and FPT gonads. (D-F”) Immunofluorescence of *vasa* and *β-catenin* of MPT, FPT with aromatase inhibition and FPT gonads. Scale bars: 50 μm.

**Table S1.**
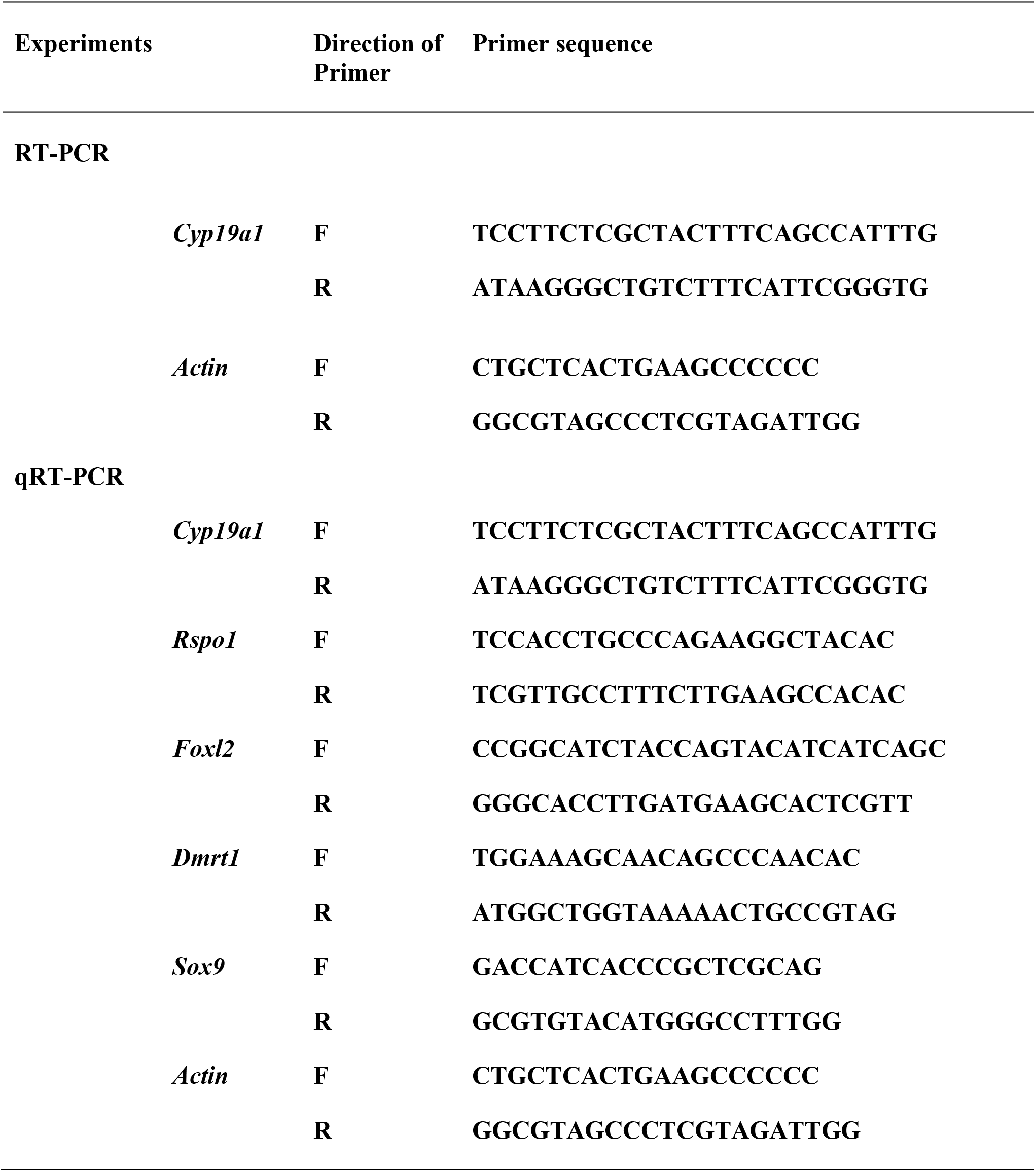
Primer list.

